# On reciprocal causation in the evolutionary process

**DOI:** 10.1101/122457

**Authors:** Erik I. Svensson

## Abstract

Recent calls for a revision of standard evolutionary theory (SET) are based in part on arguments about the reciprocal causation. Reciprocal causation means that cause-effect relationships are obscured, as a cause could later become an effect and *vice versa*. Such dynamic cause-effect relationships raise questions about the distinction between proximate and ultimate causes, as originally formulated by Ernst Mayr. They have also motivated some biologists and philosophers to argue for an Extended Evolutionary Synthesis (EES). The EES will supposedly expand the scope of the Modern Synthesis (MS) and Standard Evolutionary Theory (SET), which has been characterized as gene-centred, relying primarily on natural selection and largely neglecting reciprocal causation. I critically examine these claims, with a special focus on the last conjecture and conclude – on the contrary– that reciprocal causation has long been recognized as important both in SET and in the MS tradition, although it remains underexplored. Numerous empirical examples of reciprocal causation in the form of positive and negative feedbacks are now well known from both natural and laboratory systems. Reciprocal causation have also been explicitly incorporated in mathematical models of coevolutionary arms races, frequency-dependent selection, eco-evolutionary dynamics and sexual selection. Such dynamic feedbacks were already recognized by Richard Levins and Richard Lewontin, well before the recent call for an EES. Reciprocal causation and dynamic feedbacks is one of the few contributions of dialectical thinking and Marxist philosophy in evolutionary theory, and should be recognized as such. I discuss some promising empirical and analytical tools to study reciprocal causation and the implications for the EES. While reciprocal causation have helped us to understand many evolutionary processes, I caution against uncritical extension of dialectics towards heredity and constructive development, particularly if such extensions involves attempts to restore Lamarckian or “soft inheritance”.

## Introduction

Last year – in January 2016 – the great population biologist Richard Levins passed away (Mehta 2016). Scientifically speaking, Levins was mainly known for his pioneering models about the evolution of genetic variation and adaptive plasticity in changing environments (Levins 1968). But Levins was also a political activist, life-long committed communist and Marxist (Maynard Smith 1988). Together with his political and scientific ally at Harvard University – population geneticist Richard Lewontin – Levins published a partly controversial book in 1985 entitled *The Dialectical Biologist* (Levins and Lewontin 1985). In this book, Levins and Lewontin advocated the use of the dialectical method – as developed by German socialists and philosophers Karl Marx and Friedrich Engels – and they then applied such dialectical thinking to various problems in ecology and evolutionary biology (Levins and Lewontin 1985). Levins and Lewontin paid particular tribute to Friedrich Engels, who was interested in natural science and the new and emerging field of evolutionary biology. Engels’ book *The Dialectics of Nature* (1883) contains a series of partly unfinished essays about how dialectical thinking could help to understand the natural world. In this book, Engels explained the dialectical principle of the “transformation of quantity in to quality and *vice versa*” and illustrated this principle with an analogy of how water changes from a liquid state in to a gas as temperature increases. In terms of evolution of our own species (*Homo sapiens*), Engels argued that the human brain and hand co-evolved and influenced each other’s evolutionary trajectories through selective feedbacks, so that a larger brain made it possible to evolve more fine-scale movements of hands and fingers and *vice versa* (Engels 1883).

Although the two Marxists Levins and Lewontin were highly critical of what they called “Cartesian reductionism” which they argued dominated Western science in general and evolutionary biology in particular, they did not deny the success of this traditional research approach. However, they suggested, drawing their inspiration from primarily Engels early work, that the dialectical method should complement the traditional reductionist approach (Levins and Lewontin 1985). Given that this is actually a fairly modest claim for the justification of the dialectical method, it is somewhat surprising that their book was initially met with such an extreme hostility from a large part of the community of evolutionary biologists.

One notable exception to this negative reception of their book was Levins and Lewontin’s friend and colleague, the British evolutionary biologist John Maynard Smith, who published a review of the book in *London Review of Books* (Maynard Smith 1988). Maynard Smith was himself a former member of the Communist Party of Great Britain (CPGP) and he had thus a background in Marxist philosophy. In his critical review, Maynard Smith did not hesitate to admit that Levins and Lewontin’s dialectical method had some scientific utility (Maynard Smith 1988). He also praised Levins work as one of the best examples of how dialectical thinking could provide scientific insights about ecological phenomena and evolutionary processes beyond Cartesian reductionism (Maynard Smith 1988). Maynard Smith’s largely sympathetic although not entirely uncritical review contrasts sharply with the hostility against the book and its authors from other parts of the evolutionary biology community in the US. In retrospect and with a knowledge of subsequent history – most notably the fall of the Berlin Wall in 1989 and the collapse of the Soviet Union in 1991 – the extremely negative reactions to *The Dialectical Biologist* could perhaps be interpreted as an effect of the general political climate in the US and the ongoing Cold War. Also, many evolutionary biologists in the US were probably not aware about the crucial difference between Stalinism as an official state ideology in the Soviet Union and Eastern Europe, and the more critical intellectual Marxist analytical tradition in Western Europe.

One of the most famous chapters of *The Dialectical Biologist* has the title “The Organism as the Subject and Object of Evolution”. In this chapter, the authors build upon some earlier foundational work by Lewontin (Lewontin 1983) and use some general coupled differential equations to explore the relationship between organism (***O***) and environment (***E***). They show that organisms are not only passive *objects* of the external environment which suffer from the force of natural selection, but the organisms are also active *subjects*, who actively modify their environments, often towards their own advantage. Thus, a fit between ***O*** and ***E*** can in principle be achieved in two different ways (although not mutually exclusive); either natural selection modifies ***O*** to fit ***E***, or ***O*** modifies ***E*** to its own advantage (Okasha 2005). One empirical example of this is thermoregulation behaviours in ectothermic animals like reptiles and insects. In *Anolis*-lizards, for instance, it has been shown that because of adaptive behavioural thermoregulation, animals can “buffer” themselves against harsh thermal environments (e. g. too cold environments) by actively searching for warmer places, thereby partly counteracting selection for improved thermal physiology (Huey et al. 2003). The main point is that there is a reciprocal feedback between ***O*** and ***E***: ***E*** influences ***O*** through the process of natural selection, but ***O*** can also influence ***E*** through niche construction (Odling-Smee et al. 2003; Okasha 2005). For the consistence of terminology I shall call such feedbacks between ***O*** and ***E*** for *reciprocal causation* for the rest of this article, following the terminology by Laland and colleagues (Laland et al. 2011), although I note that Laland (2004) has also called this “cyclical causation” in one of his previous papers (Laland 2004; Dawkins 2004).

Naturalists and field biologists have long been aware that organisms are not only passive objects of selection, but can modify their environments or use adaptive habitat selection to maximize fitness (Huey et al. 2003), so in that sense Levin’s and Lewontin’s main contribution was to highlight what many already knew, and thereby encourage further investigation of these phenomena. However, it took a couple of more decades until Levin’s and Lewontin’s ideas attracted more interest from modellers. In 2003, Odling-Smee, Laland and Feldman published a book called *Niche Construction – the neglected process in evolution* (Odling-Smee et al. 2003). Building on the original foundations laid by Levins and Lewontin, they further developed the mathematical models of coupled differential equations between ***O*** and ***E***, and argued that niche construction deserved increased attention from evolutionary biologists, as it should be considered an evolutionary process, one potentially of equal importance as natural selection. While many evolutionary biologists would probably agree that niche construction and phenomena associated with reciprocal causation are interesting and important, Odling-Smee et al. (2003) were also criticized for overstretching the domain of niche construction (Brodie 2005) and their book generated considerable discussion about definitions and domains of this process (Dawkins 2004; Okasha 2005). Interestingly, Odling-Smee et al. (2003) cite Levins and Lewontin (1983) at only one page in the beginning of their volume, and neither of the terms “dialectics” or “Marxism” appear in their index. This is an interesting omission, considering the intellectual and scientific roots of niche construction and the crucial contributions by Levins and Lewontin. It is as if the Cold War was still ongoing in 2003, when *Niche Construction* was published. This might appear unfortunate, as even the otherwise skeptical John Maynard Smith was not afraid of admitting the fruitful contribution of some aspects of Marxist philosophy to evolutionary theory (Maynard Smith 1988; Maynard Smith 2001). Interestingly, two of the authors of *Niche Construction* claim to have taken the advice of Richard Lewontin, who was concerned that the use of the term ‘dialectic’ would lead to their scientific arguments being disregarded as politically motivated (Laland and Odling-Smee, personal communication). Nonetheless, niche construction theory can be viewed as implicitly embracing the dialectical method, by framing itself as a counterpoint to mainstream evolutionary biology, with the motive of stimulating research on the topic (Odling-Smee et al. 2003).

Niche construction and reciprocal causation have recently been used as arguments in calls for an Extended Evolutionary Synthesis (EES), to complement and extend the Modern Synthesis (MS), sometimes also called Standard Theory (SET)(Laland et al. 2015). These calls have, however, also met several criticisms (Welch 2016; Gupta et al 2017). Among the criticisms that have been raised against the EES are that reciprocal causation is already well-recognized in several subfields of evolutionary biology, that it is already incorporated in standard evolutionary theory and that so-called “soft inheritance” is unlikely to be important in evolution (Brodie 2005; Dickins and Rahman 2012; Welch 2016; Gupta et al 2017).

Here, I discuss this further with a focus on the role of reciprocal causation in the evolutionary process. I show that reciprocal causation features commonly in both empirical investigations and in theoretical models of both ecology and evolution, although it is seldom explicitly framed as such or couched in terms either reciprocal causation, dialectics, niche construction or the EES. Many evolutionary biologists have already implicitly or explicitly accepted reciprocal causation and unconsciously use dialectical thinking in their research practice, which calls in to question the need for urgent reform of SET and a major conceptual revision, requested by proponents of the EES. The main challenge is therefore mainly empirical rather than concepetual; namely to use existing analytical, statistical and mathematical tools to analyze reciprocal causation and spread knowledge about these tools to other subfields. I therefore suggest developing and exploiting these tools rather than calling for a major revision of evolutionary theory is a more constructive way to move research forward in these areas..

### Reciprocal causation: frequency-dependence, eco-evolutionary dynamics and co-evolution

Brodie (2005) in his review of *Niche construction* criticized the Odling-Smee et al. (2003) for painting a biased and misleading view of how evolutionary biologists study selection and its consequences:

> “The authors work hard to convince the reader that niche construction is a new ‘‘extended theory of evolution’’ that is a ‘‘co-contributor, with natural selection, to the evolutionary process itself’’ (p. 370). This argument is based on the somewhat disingenuous contention that evolutionary biologists view natural selection as an abiotic entity that is not influenced or changed by living organisms, and that ‘‘adaptation is conventionally seen as a process by which natural selection shapes organisms to fipre-established environmental ‘templates’’’ (Laland et al. 2004). This straw man is weakened by the long list of similar ideas that the authors themselves describe, from frequency-dependent selection, to coevolution, to cultural inheritance, to maternal effects. Each of these ideas (and many others) points to a general appreciation that selection is a dynamic process that changes as organisms evolve and interact with their environments. The basic tenets of niche construction can be traced back at least as far as Fisher (1930). The oft-misunderstood fundamental theorem apparently included the assumption that growing populations are expected to degrade their environments so that the positive effects of genetic increases in fitness combine with negative feedback on environmental variation for fitness (Frank and Slatkin 1992). The net result in Fisher’s view was that selection for increased fitness would not lead to any observable change in population mean fitness because evolving organisms modify their environments. The more active sense of engineering an organism’s own selection was captured early on by Mayr’s (1963) notion that behavior leads the evolution of morphology, ecology, and species differences. Through behavioral plasticity, organisms might shift niches, change diets, and move to new habitats, thereby changing selection so that ‘‘other adaptations to the new niche… are acquired secondarily’’ (Mayr 1963, p. 604). The basic premise that organisms interact with selection through a dual-direction causal arrow is not particularly novel or earth-shattering.”

From the perspective of an empirical field-oriented evolutionary biologist, I very much agree with Brodie’s characterization of the SET and the MS above, although advocates of niche construction theory would counter that while the fact that organisms modify their environments have been widely recognized, SET does not explicitly recognize this organismal agency as a direct cause of evolutionary change (Odling-Smee et al 2003). For another recent and extensive criticism of the EES and niche construction theory with additional empirical examples, see Gupta et al. (2015). Gupta et al. (2015) reviewed parts of the extensive literature of density-dependent selection, and argued that many previous empirical studies on density-dependent selection already cover many of phenomena that Odling-Smee et al. (2003) claimed are missing from the MS and SET. Central to this debate is how widely recognized is reciprocal causation in SET and among evolutionary biologists? Here, I discuss this by focusing on some phenomena which all exemplify reciprocal causation.

First, negative frequency-dependent selection (NFDS) is a well-recognized evolutionary process that involves reciprocal causation, and which was known already by the founding fathers of the MS and the early mathematical population geneticists. NFDS was explicitly incorporated in Fisher’s model for sex ratio evolution (Fisher 1930) and investigated in depth by Sewall Wright in terms of its role in maintaining genetic polymorphisms (Wright 1969). Later, NFDS became popular also in behavioural ecology, through evolutionary game theory (Maynard Smith 1982). Empirically, NFDS has been identified and studied in several field and laboratory systems, and it is a dynamic research field that has grown out of SET (Sinervo and Lively 1996; Sinervo et al. 2000; Svensson et al. 2005; Neff and Svensson 2013; Zhang et al. 2013; Le Rouzic et al. 2015). The importance of NFDS is by no means restricted to its role in maintaining genetic polymorphisms within local populations, but it can also affect population performance such as stability, productivity or extinction risk (Takahashi et al. 2014). Negative frequency-dependence might also be an important process in community ecology, where it can preserve biodiversity through rare-species advantages (Wills et al. 2006). NFDS is an example of a negative feedback loop in which a genotype’s fitness is negatively regulated by its own frequency (Fig. 1A). Agenotype can thus be said to “construct” its local selective environment (Brandon 1990; Fig. 1A). Therefore, NFDS is a prime example of a process of reciprocal causation, and it has been long been recognized as important in SET. Finally, Fisher’s Fundamental Theorem (as mentioned by Brodie 2005) does also contain a strong element of negative frequency-dependence and density-dependence through the effects of the deterioration of the environment that follows after an efficient, aggressive or highly exploitative genotype have spread in a local population and starts to encounter and interact increasingly with itself (Frank and Slatkin 1992).

**Figure 1.**
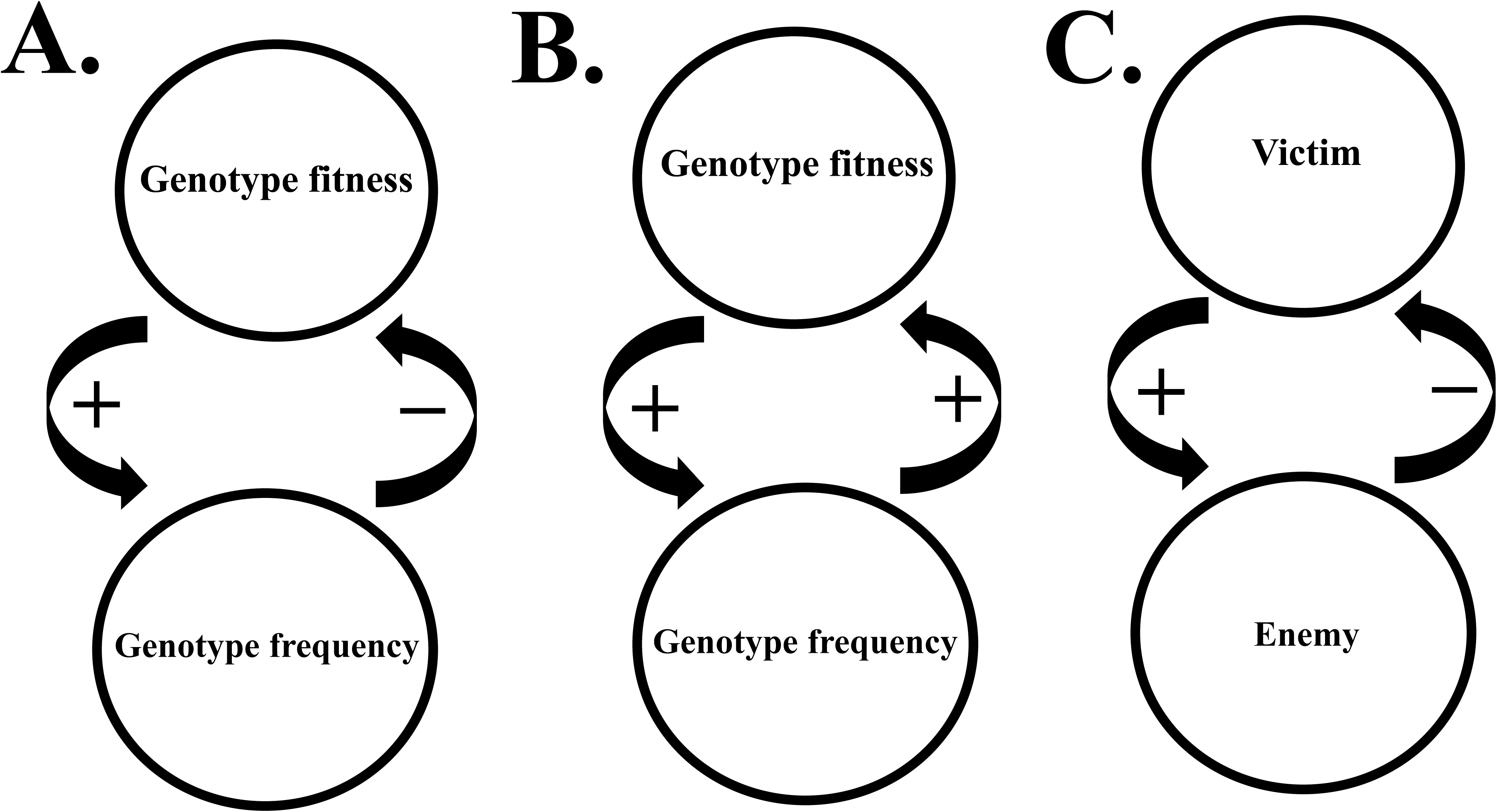
Three examples of reciprocal causation and feedbacks in the evolutionary process. **A.** Negative frequency-dependent selection (NFDS), exemplifies negative (regulatory) feedback between genotype frequency and fitness. As a genotype increases in frequency, its fitness declines, leading to the preservation of genetic diversity and genetic polymorphisms locally. The genotype thus “constructs” its own selective environment by regulating its own fitness, and the selection coefficient on the genotype changes dynamically with changing frequency. **B.** Positive frequency-dependent selection (PFDS), exemplifies positive feedback between a genotype’s frequency and its fitness, leading to fixation of the most common phenotype. **C.** Enemy-victim coevolution is an example of a negative feedback, that can either lead to stable equilibria or co-evolutonary cycles (e. g. “Red Queen” evolutionary dynamics). The enemy and the victim can belong to different species (e. g. predators or prey, parasites or hosts) or the same species (e. g. males and females).

However, also positive frequency-dependent selection (PFDS) is an important evolutionary process that exemplifies reciprocal causality (Fig. 1B). Under PFDS, a genotype’s fitness will increase as it becomes more common. PFDS leads to the loss of polymorphism and the fixation of locally common genotypes, in contrast to NFDS which plays a conservative role in population divergence (Svensson et al. 2005; cf. Fig. 1A vs. B). One classical example where PFDS plays an important role in evolution is Müllerian mimicry in *Heliconus*-butterflies, where locally common warning colouration patterns are favoured due to predator learning (Chouteau et al. 2016). Another example comes from classical models of sexual selection through female choice, in particular the Fisher-Lande-Kirkpatrick model (Fisher 1930; Lande 1981; Kirkpatrick 1982; Prum 2010). In this model, there is genetic variation in both female preferences for a male secondary sexual trait and the male trait itself. Female choice and non-random mating leads to the buildup of linkage disequilibrium (LD) between female preference alleles and male trait alleles, and a genetic correlation between these disparate traits forms, even if the traits are governed by separate sets of loci (Kirkpatrick 1982). Provided that this genetic correlation becomes of sufficiently high magnitude, a tipping point might be reached and a “runaway” process can get started whereby PFDS drives male-expressed trait alleles and the corresponding female preference alleles to fixation (Prum 2010). The important message here is that selection becomes self-reinforcing: given set of preferences, male-expressed trait-alleles spread through the synergistic effects between PFDS and the LD that was generated by the female preference. Thus, not only do the traits of males and females evolve, but so does the selective environment itself – an excellent example of reciprocal causation and feedback between organism and environment (Levins and Lewontin 1985). Recently, positive frequency-dependence has also been suggested to be important in maintaining diversity of ecologically equivalent species (e. g. those formed by sexual selection alone) on regional scales (M’Gonigle et al. 2012), and it might also play an important role in community ecology through priority effects (De Meester et al. 2016). Even more generally, positive feedbacks and runaway processes have been suggested to be important in human social evolution in coalition formation and cooperative networks (Gavrilets et al. 2008) and in ecosystem ecology and climate science (Scheffer et al. 2001; Malm 2007).

Reciprocal causation has also a key role in the field of “eco-evolutionary dynamics” (Schoener 2011; Hendry 2016), where the focus are the bidirectional feedbacks between ecological (e. g. population dynamics) and evolutionary processes (e. g. genetic change within populations). Eco-evolutionary dynamics is expected when ecological and evolutionary time scales converge, such as in the case of rapid evolution, e. g. due to human-induced environmental changes (Hendry et al. 2017). Under such scenarios does not only ecological change affect genetic change, but also *vice versa*: genetic changes can feed back in to ecology and influence population dynamics (Sinervo et al. 2000; Yoshida et al. 2003). Note that this breakdown of the separation between ecological and evolutionary time scales can be viewed as a problem for certain modelling approaches, such as Adaptive Dynamics (AD), where a strict separation between ecological and evolutionary processes is a core assumption (Dieckmann and Doebeli 1999; Waxman and Gavrilets 2005). Such dynamic feedbacks between ecology and evolution will without doubt continue to be explored in the future, and they show that reciprocal causation forms a key part of a rapidly growing research field that has largely developed independently from niche construction theory.

Interestingly, Schoener (2011) called eco-evolutionary dynamics for “the newest synthesis”between ecology and evolution. Schoener’s call for this new synthesis was independent of Pigliucci, Laland and colleagues (Pigliucci 2007; Pigliucci and Müller 2010; Laland et al. 2015). From a strict empirical viewpoint, eco-evolutionary dynamics have contributed more towards the development of a concrete empirical research program in a shorter time than has the EES sofar, although it is only recently that the latter has tried to formulate an empirical research program (Laland et al. 2015). I also agree with Welch (2016) that there is an unfortunate tendency in evolutionary biology to repeatedly use the richness of understudied and underappreciated phenomena and processes in our field as arguments for the urgent need of reform and to proclaim the arrival of new syntheses. It is worth emphasizing that there are also several other, more or less parallel attempts to call for new evolutionary synthesis, which are either based on conceptual considerations (Arnold 2014) or on new methods (Losos et al. 2013). These other synthesis-attempts are more modest in their scope than the EES, the latter which embraces an explicit counterpoint to SET, i. e. a dialectical approach.

Reciprocal causation also forms a key component in studies of co-evolution, either within or between species, such as enemy-victim interactions (Fig. 1C). For instance, under interlocus contest evolution (ICE) between male and female over mating rates (i. e. sexual conflict), males evolve traits that increase their mating success, whereas females evolve resistance towards excessive male mating harassment (Rice and Holland 1997). Under certain conditions, such antagonistic male-female sexual interactions can lead to intraspecific co-evolutionary arms races (Rice 1996) and even speciation (Gavrilets 2000). Females can also evolve either resistance or tolerance to male mating harassment (Arnqvist and Rowe 2005; Gosden and Svensson 2009; Svensson and Raberg 2010), and such sexual antagonism can also, as an alternative outcome, result in the evolution of male and female genetic clustering or polymorphisms (Gavrilets and Waxman 2002; Svensson et al. 2009; Karlsson et al. 2013; Karlsson et al. 2014; Le Rouzic et al. 2015). These antagonistic male-female interactions sometimes result in “Red Queen”-dynamics and co-evolutionary feedbacks between male and female fitness interests (Rice and Holland 1997). This exemplifies how reciprocal causation is deeply embedded in the empirical research traditions of evolutionary genetics and evolutionary ecology. Reciprocal causation also embedded in the idea of geographic coevolutionary selection mosaics across the landscape of interacting species (Gomulkiewicz et al. 2000; Nuismer et al. 2000; Thompson 2005). As in the case of frequency-dependent selection and eco-evolutionary dynamics, appreciation of reciprocal causation is the norm rather than the exception among many active empiricists in evolutionary biology.

Reciprocal causation has also been explicitly incorporated in models and in empirical investigations in the evolutionary quantitative genetics tradition, where researchers are using the statistical methods that were developed by Russel Lande and Stevan Arnold (Lande 1976; Lande and Arnold 1983; Endler 1986). Examples of such studies incorporating reciprocal causality include indirect genetic effects in social evolution (Moore et al. 1998; Wolf et al. 2001), the evolutionary dynamics of maternal effects (Kirkpatrick and Lande 1989; Wade 1998) and analyses of how interspecific interactions shape selection pressures on phenotypic traits (Ridenhour 2005). Empirical selection studies nowadays are not only aiming to quantify selection differentials and selection gradients, but researchers actively strive to understand the ecological causes of selection, whether those causes are predators, intra-or interspecific competitors (Wade and Kalisz 1990; Svensson and Sinervo 2000; Svensson and Friberg 2007; Calsbeek and Smith 2007; Calsbeek et al. 2012; Kuchta and Svensson 2014). It is a very long time ago since evolutionary ecologists were simply satisfied by having quantified selection (Lande and Arnold 1983; Wade and Kalisz 1990). Nowadays, evolutionary ecologists are busy understanding the ecological causes of selection (Siepielski et al. 2017) and few journals in evolutionary biology publish studies where selection coefficients are simply presented without any ecological context (and rightly so).

This brief review illustrates how positive and negative feedbacks are a fact of life. Here, I find myself largely in agreement with advocates of the EES in emphasizing the ubiquity of reciprocal causation in evolution, but equally in agreement with critics wo maintain that reciprocal causation is already well recognized within SET.

### Analytical and empirical tools for studying reciprocal causation

As I have discussed and exemplified above, reciprocal causation is hardly controversial among evolutionary biologists and widely recognized in several subfields in evolutionary biology. Moreover, reciprocal causality has been recognized as important for several decades and well before the formalization of niche construction (Odling-Smee et al. 2003) and more recent calls for an EES (Laland et al. 2015). Few empirical and theoretical evolutionary biologists today adhere to a simple unidirectional causality. Even Ernst Mayr himself expressed a more dynamical view of causality in other contexts and publications than he did in his distinction between proximate and ultimate causes (Laland et al. 2011). Mayr’s views of the role of behaviour as a “pace maker” in evolution (Mayr 1963), strikes me as being much more sophisticated than the picture of unidirectional causality that has been described by Laland et al (2011). Mayr’s view of a crucial role of behaviour in the evolutionary process is clearly compatible with feedbacks between the organism and its environment. Mayr’s surprisingly early insights on the issue has clear similarities with similar views expressed several decades later by West-Eberhard, Levins and Lewontin (West-Eberhard 1983; Levins and Lewontin 1985), albeit not developed in detail by him.

If reciprocal causation is then so widely recognized – at least in several key fields – why then is it not more studied? Here I question the claim that there is a major conceptual barrier to recognize reciprocal causation, as maintained by the architects of niche construction theory and the EES (Odling-Smee et al. 2003; Laland et al 2011; 2015). Rather, the answer is probably that there are enormous logistical and empirical challenges, and not all researchers are aware of suitable analytical tools. Progress in the field of evolutionary biology is perhaps more often limited to methods these days than to lack of conceptual insights. It therefore becomes more urgent to communicate between subfields so that researchers become aware of which analytical and empirical tools that are aready available, but which are underutilized. I therefore agree fully with Laland et al. (Laland et al. 2013) that different subfields in biology should become better integrated. However, I doubt that such integration will need or necessarily be facilitated by the adoption of an EES, at least not in its current rather vague form. Rather, the main motivation for fostering integration between different fields in biology is that statistical, mathematical and other analytical tools suitable for studying reciprocal causation are underutilized in some subfields, and scientific communication would facilitate their spread.

One such tool that is clearly underutilized in many areas of evolutionary biology and which is excellently suited to analyze direct and indirect effects is path analysis and structural equation modelling (SEM)(Shipley 2002; see also Laland et al 2011). Although path analysis has been advocated as a suitable tool in selection analyses on phenotypic traits (Kingsolver and Schemske 1991), path analyses of selection are still relatively few (Sinervo and DeNardo 1996). This is unfortunate, as path analyses and SEM are powerful tools to incorporate information about how the development and expression of phenotypic traits are influenced by local social, biotic and abiotic environments, and how traits in turn affect fitness and thereby are linked to selective environments (Svensson et al. 2001; Gosden and Svensson 2009). Moreover, path analysis can also be combined with experimental manipulations – either of phenotypic traits, of local selective environments, or both (Sinervo and Basolo 1996; Svensson and Sinervo 2000). Integrative studies combining path analysis, analysis of causation and experimental manipulations will increase our knowledge about organism-environment feedbacks and therole of such feedbacks in the evolutionary process (Svensson et al. 2002). Empirical information from covariance or correlation matrices can be translated in to causal quantitative models, whereby SEM provides a powerful tool to evaluate the fit of various alternative models (Shipley 2002).

Another underutilized tool to study reciprocal causation in the evolutionary process is time-series analysis (Le Rouzic et al. 2015). Time-series analysis has perhaps been more used by ecologists interested in population dynamics than by evolutionary biologists, but it holds great promise as a tool to infer the processes driving ecological and genetic dynamics of interacting genotypes within species (Moorcroft et al. 1996; Pemberton et al. 1998; Sinervo et al. 2000; Le Rouzic et al. 2015) or in analyses of interspecific interactions (Yoshida et al. 2003). Time-series analysis could be especially powerful if it would be combined with experimental manipulations of putative causal ecological agents of selection (Wade and Kalisz 1990; Svensson and Sinervo 2000). I anticipate that evolutionary time-series analysis will become an important tool in future studies dealing with eco-evolutionary dynamics, intra-or interspecific co-evolutionary processes in natural populations (Le Rouzic et al. 2015; Hendry 2016).

Other promising research approaches to investigate reciprocal causation and dynamic feedbacks between organisms and their local environments include studies of non-random dispersal with respect to phenotype or genotype (Edelaar et al. 2008; Eroukhmanoff et al. 2011) and consequences for matching habitat choice (Edelaar and Bolnick 2012), quantitative studies on the dynamics of niche evolution using phylogenetic comparative methods (Wiens 2011; Wiens et al. 2011), and experimental field studies on how animals use regulatory behaviours to maintain physiological homeostasis (Huey et al. 2003). Taken together, there is therefore a rich diversity of powerful empirical and analytical tools available to evolutionary biologists who are seriously interested in understanding how reciprocal causation and dynamic feedbacks between ecological and evolutionary processes influence organisms, from individuals to populations, species and higher taxa.

### A cautionary note about extending reciprocal causation to heredity and constructive development

As I have shown in this article, reciprocal causation is already widely recognized in several fields, particularly those at the interface between ecology and evolution. Laland et al. (2015) do of course not deny the existence of such previous studies but they argue that:

> “However, reciprocal causation has generally been restricted to certain domains (largely to direct interactions between organisms), while many existing analyses of evolution, habit-or frequency-dependent selection are conducted at a level (e. g. genetic, demographic) that removes any consideration of ontogeny. Such studied do capture a core structural feature of reciprocal causation in evolution – namely, selective feedback – but typically fail to recognize that developmental processes can both initiate and co-direct evolutionary outcomes”
>
> — (p. 7. Laland et al. 2015).

Thus, Laland et al. (2015) admit that reciprocal causation is and has often been studied by evolutionary biologists, but they argue that ontogeny and development should be incorporated in such analyses. I hardly disagree here, and I think incorporating the role of development and ontogeny in studies of (say) frequency-dependent selection, eco-evolutionary dynamics, co-evolution and analyses of selection is likely to yield many novel and important insights. However, the reason that development has not been incorporated in that many previous studies in this field is not that the researchers in question rely on an outdated and simple view of unidirectional causation, as implied by Laland et al. (2015). The reason is more likely a practical one: it is extremely difficult and empirically challenging to understand and study reciprocal causation even at single ontogenetic level, such as among adults. I therefore disagree with Laland et al. (2011; 2013) when they imply that the lack of consideration of development in past studies is due to the lasting legacy of Ernst Mayr’s proximate-ultimate dichotomy, and their suggestion that evolutionary biologists in general implicitly adher to an outdated view of unidirectional inheritance. Rather, the lack of studies of this kind reflect legitimate and difficult empirical challenges and I am not convinced that the EES-framework alone can solve these problems, unless some more concrete novel methodological or analytical tools are provided.

Moreover, evolutionary geneticists and evolutionary ecologists have actually paid attention to the interplay between ontogeny and selection. For instance, researchers have modelled and investigated how selection pressures change both in magnitude and sign during the organism’s life cycle (Schluter et al. 1991; Barrett et al. 2008). For instance, there is much interest and ongoing theoretical and empirical research aiming to integrate and model the interaction between interlocus sexual conflict at the adult stage over the reproductive interests of males and females, with intralocus sexual conflict experienced earlier in ontogeny (Rice and Chippindale 2001; Chippindale et al. 2001; Barson et al. 2015; Pennell et al. 2016). There is also an increased appreciation of how alternative reproductive strategies shape ontogenetic trajectories, and how the same ontogenetic trajectories in turn affect adult phenotypes (Neff and Svensson 2013). Laland et al. (2015) wish to extend the domain of reciprocal causation from the interaction between ecological and evolutionary processes (as discussed in this article) to the domain of organismal development, what they call “constructive development”. I will not dwell too deeply in to this here, due to space limitations, except that I note that there is of course no *a priori* reason why reciprocal causation and dialectical thinking should not be possible to apply also to development. However, constructive development is also perhaps the aspect of the EES that is most controversial and which has sofar been met with most resistance. This resistance is partly understandable and justified, on historical grounds. The architects of the MS had to work hard to get rid of popular evolutionary mechanisms of inheritance and evolution that in hindsight have clearly turned out to be wrong, including orthogenesis, vitalism, saltationism and Lamarckian inheritance (Smocovitis 1996; Mayr and Provine 1998). The increasing interest in epigenetic inheritance is certainly justified and will most likely lead to new empirical insights. Clear cases of epigenetic inheritance now exists (Dias and Ressler 2014) and it is now mainly an empirical issue to understand the importance of such effects and how widespread they are. However, claims that epigenetic inheritance can play a major role in explaining macroevolutionary phenomena (Pigliucci and Murren 2003) or that such epigenetic inheritance would imply a major role for Lamarckian inheritance and saltations in evolution (Jablonka and Lamb 2005; 2008) have been criticized as speculative and without empirical footage (West-Eberhard 2007; Dickins and Rahman 2012). The view that so-called soft inheritance (as defined by Jablonka and Lamb 2008) necessitates a replacing the MS with an entirely new theoretical framework shows that the Lamarckian temptation is still present among a small minority of biologists, but I will not dwell in to this further here. For in-depth criticisms of this minority position, see West-Eberhard (2007) and (Haig (2007).

Maynard Smith (1988; 2001) who certainly admitted that Marxist philosophy and dialectical thinking could have a constructive influence on evolutionary theory cautioned against uncritical extenson of dialectics to heredity and development. Maynard Smith’s cautionary point should also be taken seriously by those today who argue for constructive development, including Laland et al. (2015). In a modest form, constructive development is entirely compatible with quantitative genetics theory, where it is explicitly recognized that gene expression is strongly environment-dependent (e. g. Lancaster et al. 2015), if such environment-dependent gene expression is heritable, and that genes, environmental conditions, gene-gene interactions (epistasis) and gene-by-environment interactions (GEI:s) all influence the development of the adult phenotype (Lynch and Walsh 1998). In a stronger form, constructive development could entail questioning the unidirectional flow of information from the genotype to the phenotype, based on (vulgar) dialectical arguments about reciprocal causality (Maynard Smith 1988; 2001). The history of genetics in the Soviet Union under Stalin’s regime and under Trofim Lysenko is a particularly interesting and tragic case in point. Under Stalinism, Engels dialectical thinking became elevated to “natural laws” and became official dogma. Stalin and Lysenko rejected Mendelian genetics on the grounds that it was “undialectical” and hence an example of “bourgeoisie science”. Lysenko instead promoted an alternative and ideological state-supported official view of constructive phenotype development based on neo-Lamarckism, where acquired characters could be inherited and transmitted genetically to future generations. Lysenko’s rejection of Mendel’s and Weisman’s views of heredity was, however, based on ideological, rather than scientific principles and hence a prime example of an ideological fallacy. The important point here is that Mendelism and unidirectional causality from heredity to phenotypes might seem undialectical, but it does not matter if this is how heredity works. After all, it must be reality that should be the guiding principle for scientific investigations, in evolutionary biology and other fields. Rejecting empirical findings in genetics solely on the basis that they seem undialectical is clearly unscientific and shows the danger of applying a philosophical framework uncritically, and erroneously elevating dialectics to the status of natural laws.

Few evolutionary biologists today would claim that the genotype-phenotype map is perfectly linear, that all genetic variation is additive and that environmental effects are unimportant in phenotype development. On the contrary, most evolutionary biologists are well aware that additive effects of genes, environmnets, GEI:s and epistasis all jointly influence phenotypic development. The ideas of cultural evolution and various forms of non-genetic inheritance (including ecological inheritance) can and does play some role in between-generation change of phenotypes is also gaining some acceptance. The role of epigenetic inheritance in evolution does not necessarily require a major revision of SET, if we recognize the crucial difference between molecular and evolutionary definitions of the term “gene” and if we treat “epialleles using the same analytical framework as classical genetic alleles in population genetics (Lu and Bourratt 2017). However, it is considerably more controversial to argue for reciprocal causation if that implies that environments could causally influence genetic inheritance, i. e. a revival of acquired genetic inheritance and neo-Lamarckism. That the the germ line is separated from the soma and that phenotypes (as far as we know) cannot causally influence heredity in an adaptive fashion may seem undialectical, but are as close to scientific facts as they could possibly be. Extremely strong evidence would be required for any claims that the Weismannian germline-soma separation is not valid anymore, that the so-called “Central Dogma” of molecular genetics does not hold up (Maynard Smith 1988; 2001) or that soft inheritance (Jablonka and Lamb 2008) is a major player in evolution (Haig 2007; Dickins and Rahman 2012). Thus, while most evolutionary biologists would have few problems in accepting reciprocal causality in ecological and evolutionary interactions between organisms and their selective environments, few are – for quite understandable historical reasons - prepared to accept reciprocal causality in constructive development and genetics, unless strong empirical evidence is presented for such claims.

### Can evolutionary quantitative genetics provide a bridge between MS and the EES?

Laland et al. (2015) reviewed and compared the structures, assumptions and predictions of the EES and contrasted these against the MS. Among the core assumptions of the MS that they identifed were “The pre-eminence of natural selection” and “Gene-centred perspective” (their Table 1). They further criticized the “blueprint”, “program” and “instruction” metaphors in genetics and the MS, and contrasted the use of these terms against their own views on constructive development. Here, I take issue with these claims, and argue that these characterizations provides a wrong, or at least very biased picture of both the current state-of-the-art of SET and the history of the MS. Laland et al. (2015) have underestimated the flexibility and scope of evolutionary genetics, a criticism that was recently developed in more detail by Gupta et al. (2015).

With respect to the claim of the pre-eminence of natural selection in the MS, it must be emphasized that most evolutionary biologists today, including many molecular population geneticists, do not agree with this claim (see also Welch 2016 for further discussion). On the contrary, leading molecular population geneticists are highly critical of what they consider an excessive adaptationist research programme in evolutionary and behavioural ecology. Many evolutionary biologists on the contrary that random processes such as genetic drift should more often be used as a null modell and point of departure, before invoking natural selection (Lynch 2007). Historically, and from the very beginning of the MS, the non-adaptive process of genetic drift was considered to have a much more powerful evolutionary role than it perhaps deserved to have, something which only became clear after extensive empirical investigations in both the field and in laboratory studies (Provine 1986).

With respect to the characterization of the MS as gene-centred, many organismal biologists, particularly those working in the evolutionary quantative genetics tradition, are likely to strongly disagree. Evolutionary quantitative genetics focus on organisms and use their phenotypic traits (variances and covariances) as its point of departure, and thereby ignores underlying molecular genetic and developmental mechanisms behind these traits (Lynch and Walsh 1998). This might be perceived as a weakness with the evolutionary quantitative genetics approach, however, it can also be perceived as a strength (Steppan et al. 2002) as quantitative genetics through this procedure become liberated from the tyranny of genetic details in classical population genetics, as argued forcefully recently by Queller (2017). Moreover, the trait variance decomposition approach in quantitative genetics would work equally well in a non-DNA world with non-genetic inheritance, as long as there is trait heritability, i. e. this mechanism-free approach is general and flexible. For instance, the Price Equation does not assume that heredity is based on DNA, but is based on the phenotypic resemblance between relatives, such as parents-offspring covariance (Frank 1995; 1997). Thus, the quantitative genetic approach does already present a substantial extension of classical population genetics that it grew out from, and could potentially be extended further to account for various forms of non-genetic inheritance, such as ecological inheritance (see Helanterä and Uller 2010 for discussion). Quantitative genetics does already partly take constructive development in to account by modelling not only additive genetic variances and covariances, but also environmental components, dominance variation, epistasis and GEI:s (Lynch and Walsh 1998). Moreover, these different variance components are not static, but they are dynamic and can evolve. For instance, after population bottlenecks, epistatic variance can be converted to additive genetic variance (Meffert et al. 2002) and models of the Fisherian Runaway process of sexual selection have revealed that genetic covariances can evolve through a dynamic feedback between the selective environment (female choice) and male secondary sexual traits (Kirkpatrick 1982). Finally, it is also worth emphasizing that natural selection can be viewed as both an ultimate and proximate explanation, as argued recentlyby Gupta et al. (2017). The process of natural selection has actually nothing to do with genetics, and questions about the causes of selection are also questions about ecological selective agents, which have their origin in the external environment (Wade and Kalisz 1990). Therefore, in this research tradition, genes are certainly not the main causal agents explaining evolution by natural selection; it is instead the selective environment that is the main causal agent (cf. Brandon 1990; Wade and Kalisz 1990).

## Conclusions

Reciprocal causation is frequent in many studies of evolutionary processes, particularly those involving interactions between organisms, both within or between species. Research on reciprocal causation has a long tradition in evolutionary biology. Reciprocal causation was studied well before the recent calls for an EES, although not explicitly under the umbrellas of niche construction, developmental plasticity and cultural evolution. Evolutionary biologists today – particularly those working at the interface between ecology and evolution seldom have the simplified view of unidirectional causation as sometimes claimed and reciprocal causation is already an essential part of the conceptual framework of many empirical biologists. Apart from the subfields I have discussed in this article, there are also several other emerging areas where reciprocal causation is deeply embedded. For instance, non-random dispersal of phenotypes and matching habitat preferences (Edelaar et al. 2008; Edelaar and Bolnick 2012; Eroukhmanoff et al. 2011) shows that organisms are not solely passive objects of evolution, but evolutionary subjects in their own right, with some degree of independence (cf. Levins and Lewontin 1985). Similarly, niche conservatism (Wiens et al. 2010) is common in many organisms, meaning that organisms actively track habitats where their fitness is maximized, rather than passively evolving *in situ*. Niche conservatism has many interesting consequences for speciation (Wiens 2004), thermal adaptation (Svensson 2012) and thermoregulatory behaviours (Huey et al. 2003).

Evolutionary biology is a rich and diverse discipline that spans many levels of biological organization and which covers many different types of questions. This diversity of our discipline is a strength, but also comes with a cost: it is relatively easy to find areas where more research would be needed and topics that have been relatively little explored (Welch 2016). The existence of such knowledge gaps is presumably the primary science-sociological explanation for why calls for major revision of evolutionary theory or attempts to formulate new synthese**s** appear with regular intervals (Welch 2016). This is disputed by advocates of the EES who maintain the push for a new perspective arises not only from knowledge gaps but when new data, theoretical findings and and approaches collectively suggest an alternative causal understanding of evolution (Laland et al. 2015). We have seen several more or less independent attempts to formulate new syntheses only during the last decade (Pigliucci 2007; Pigliucci and Müller 2010; Schoener 2011; Losos et al. 2013; Arnold 2014; Laland et al. 2015). These attempts were preceeded by other calls in the past (Gould 1980). However, as I have argued elsewhere (Svensson and Calsbeek 2012a), new syntheses do not automatically establish themselves in the evolutionary research community because some biologists think that they are warranted. Rather, new syntheses grow organically, and become established only if they provide some new analytical, experimental, mathematical or statistical tools that moves the research field forward. The MS was never such a monolithic research paradigm as sometimes portrayed by some critics (Jablonka and Lamb 2005; Laland et al. 2015). Rather, the MS was a loose, albeit largely successful research framework and attempt to unify very heterogeneous and different branches of biology (Smocovitis 1996; Mayr and Provine 1998). Some have even questioned the existence of the MS as a clearly separated and identifiable historical period, and have argued that the term synthesis should now be abandoned as it is not valid anymore (Cain 2009). The MS contained several very conflicting perspectives on evolutionary biology, both between different branches of population genetics (Provine 1986; Frank and Slatkin 1992; Coyne et al. 2000; Wade and Goodnight 1998) and between researchers focusing on micro- vs. macroevolution (Eldredge and Gould 1972; Charlesworth et al. 1982; Futuyma 2015). Past and ongoing debates about selection versus neutralism in explaining genetic variation (Lewontin 1974; Gillespie 1991) and the role of population structure, genetic drift and mass-selection in large panmictic populations (Coyne et al. 2000; Wade and Goodnight 1998) all illustrate that the MS has been continually evolving and adapting, from a flexible minimum platform that has survived several replacement attempts (Smocovitis 1996; Svensson and Calsbeek 2012b). The MS will therefore most likely probably continue to evolve and slowly adapt also in the future (Arnold 2014). Modern evolutionary biology and SET has also already moved considerably beyond the original scope of MS. In fact, it can be argued that substantial extensions of the MS already took place several decades ago, e. g. with the incorporation of neutral theory and the development of evolutionary quantitative genetic theory and methodology, complementing the classical population genetic tradition (Queller 2017).

The perspective put forward in this article is largely an empiricist one. While I disagree with Laland et al. (2011;2015) that reciprocal causality is that neglected in evolutionary biology, I fully agree with them that it should become more widely appreciated and studied. The study of reciprocal causation need to move beyond rhetoric and position papers and become operational. The insights that organisms construct their own environments and appreciation of organism-environment feedbacks as well as the discussion whether there exist empty niches (or not) are all interesting but they need to be translated to a rigorous empirical research program, for instance, by utilizing some of the analytical tools I have discussed in this article. It remains to be seen if and how the EES can be translated in to a productive research program, but an attempt to do this is now underway (http://extendedevolutionarysynthesis.com/).The widespread existence of reciprocal causation should not be taken as an argument that cause-effect relationships are empirically impossible to study, but should rather motivate researchers to dissect long causal chains in to smaller operational study units to better understand the evolutionary process. A system of temporally separated factors of reciprocal causation can always be broken down in to separate linear causal links in a longer chain of events to facilitate understanding and analysis. Of course, we need to appreciate the crucial difference between the ecological and selective environment (Levins and Lewontin 1985; Brandon 1990), but this conceptual challenge should not hinder the development of operational research tools in empirical studies.

Reciprocal causation is already deeply embedded in many – perhaps the majority – of evolutionary processes, and should therefore be a natural and major research focus. Although broader appreciation of the role of reciprocal causation is unlikely to lead to a new paradigm shift (Laland et al. 2013) and is probably not a sufficient reason to call for major revision of evolutionary theory (Laland et al. 2015), reciprocal causation is nevertheless a good example of how Marxist philosophy and dialectical thinking have had a positive influence on the development of our field. Early insights about reciprocal causation can of course also be traced from other research traditions than Engels dialectical methods, such as from cybernetics (Wiener 1948) and from cyclical causal dependencies in early predator-prey models (Lotka 1910; Volterra 1926). We should nevertheless not hesitate to embrace the concept of reciprocal causality and acknowledge the contributions of Levins and Lewontin and the dialectical method (Levins and Lewontin 1985). Engel’s surprisingly early insights and his dialectical method can – if they are applied critically as a method rather than being treated as natural laws – still provide important insights to understand evolutionary processes. For instance, the dialectical principle of the transformation of quantity in to quality can be understood as an early insight by Engels of phase transitions, non-linear changes, hysteresis, critical thresholds and tipping points and rapid (non-gradual) switches between alternative states in ecology and evolution. Such ideas have been successfully incorporated in models of human social evolution (Carneiro 2000; Gavrilets et al. 2008), reproductive isolation and speciation (Gavrilets and Gravner 1997; Nosil et al. 2017) and in ecosystem ecology (Scheffer et al. 2001). Likewise, it is tempting to interpret Maynard Smith’s interest later in life for major evolutionary transitions (Maynard Smith and Szathmary 1988) at least as partly influenced by his background in Marxist philosophy and appreciation of dialectics, as this is an excellent example of the transformation of quantity in to quality in evolutionary biology.

## Acknowledgements

Funding for my research has been provided by the Swedish Research Council (VR: grant no.: 621-2012-3768). Many of the ideas put forward in this paper have grown out of dialectical scientific discussions with Tobias Uller. I would also like to thank Charlie Cornwallis, Kevin Laland, John Odling-Smee and John Welch for comments and input at various stages of manuscript preparation. The author is part of a recent international research consortium entitled *Putting the Extended Evolutionary Synthesis to Test*, funded by John Templeton Foundation (main PI:s Kevin Laland and Tobias Uller). Although the views put forward in this paper are somewhat critical to the EES as a concept, the author wish to thank those who think otherwise for unconsciously stimulating me to write this paper and summarize my thoughts on the topic. Evolutionary biology is a vibrant discipline with room for many different and divergent viewpoints. It is my hope that this paper will contribute to an important and ongoing discussion about the future our research field.

